# Novel functional insights revealed by distinct protein-protein interactions of the residual SWI/SNF complex in SMARCA4-deficient small cell carcinoma of the ovary, hypercalcemic type

**DOI:** 10.1101/794776

**Authors:** Elizabeth A. Raupach, Krystine Garcia-Mansfield, Ritin Sharma, Apurva M. Hegde, Victoria David-Dirgo, Yemin Wang, Chae Young Shin, Lan V. Tao, Salvatore J. Facista, Rayvon Moore, Jessica D. Lang, Victoria L. Zismann, Krystal A. Orlando, Monique Spillman, Anthony N. Karnezis, Lynda B. Bennett, David G. Huntsman, Jeffrey M. Trent, William P. D. Hendricks, Bernard E. Weissman, Patrick Pirrotte

**Affiliations:** Integrated Cancer Genomics, Translational Genomics Research Institute, Phoenix, AZ, USA; Collaborative Center for Translational Mass Spectrometry, Translational Genomics Research Institute, Phoenix, AZ, USA; Department of Pathology and Laboratory Medicine, University of British Columbia, Vancouver, BC, Canada; Department of Molecular Oncology, British Columbia Cancer Research Centre, Vancouver, BC, Canada; Lineberger Comprehensive Cancer Center, University of North Carolina, Chapel Hill, NC, USA; Department of Pathology and Laboratory Medicine, University of North Carolina, Chapel Hill, NC, USA; Gynecologic Oncology, Obstetrics, & Gynecology, Baylor Scott & White Health, Dallas, TX, USA; Department of Pathology and Laboratory Medicine, University of California Davis Medical Center, Sacramento, CA, USA; Center for Autophagy Research, University of Texas Southwestern Medical Center, Dallas, TX, USA; Department of Obstetrics and Gynaecology, University of British Columbia, Vancouver, BC, Canada

## Abstract

Chromatin remodeling plays a critical role in tumor suppression as demonstrated by 20% of human cancers bearing inactivating mutations in SWI/SNF chromatin remodeling complex members. Mutations in different SWI/SNF subunits drive a variety of adult and pediatric tumor types, including non-small cell lung cancers, rhabdoid tumors, medulloblastomas, and ovarian cancers. Small cell carcinoma of the ovary hypercalcemic type (SCCOHT) is an aggressive subtype of ovarian cancer occurring in young women. Nearly all (>98%) SCCOHTs have inactivating mutations in *SMARCA4*, which encodes 1 of 2 mutually exclusive catalytic subunits of the SWI/SNF complex. Less than half of SCCOHT patients survive 5 years despite aggressive surgery and multimodal chemotherapy. Empirical support for effective SCCOHT treatments is scarce, in part because of the poor understanding of SCCOHT tumorigenesis. To gain insight into the functional consequences of SWI/SNF subunit loss, we defined SWI/SNF composition and its protein-protein interactions (PPIs) by immunoprecipitation and mass spectrometry (IP-MS) of SWI/SNF subunits in 3 SCCOHT cell lines. Comparing these results to a cell line containing a wild-type SWI/SNF complex, the interaction of most canonical core SWI/SNF subunits was observed in all SCCOHT cell lines at a lower abundance. The SCCOHT SWI/SNF also lacked ATPase module subunits and showed a drastic reduction in PBAF-specific subunit interactions. The wild-type and SCCOHT SWI/SNF subunits immunoprecipitated a shared set of 26 proteins, including core SWI/SNF subunits and RNA processing proteins. We observed 131 proteins exclusively interacting with the wild-type SWI/SNF complex including isoform-specific SWI/SNF subunits, members of the NuRD complex, and members of the MLL3/4 complex. We observed 60 PPIs exclusive to the SCCOHT residual SWI/SNF shared in at least 2 of the 3 SCCOHT cell lines, including many proteins involved in RNA processing. Differential interactions with the residual SWI/SNF complex in SCCOHT may further elucidate altered functional consequences of SMARCA4 mutations in these tumors as well as identify synthetic lethal targets that translate to other SWI/SNF-deficient tumors.

## Introduction

Nearly 20% of human cancers bear mutations in the switch/sucrose non-fermenting (SWI/SNF) chromatin remodeling complex^1,2^, ranking it among the most highly mutated protein complexes in cancers. The SWI/SNF complex, first characterized in yeast, is best known for its ability to activate gene expression by facilitating formation of nucleosome-depleted regions at gene promoters^3–7^. The mammalian SWI/SNF complex has important roles in development and consists of at least 3 major isoforms: BRG1(SMARCA4)-associated factor (BAF), poly-bromo-BAF (PBAF), and non-canonical BAF (ncBAF), which can either activate or repress gene expression in different contexts^8–12^. These isoforms share a set of core subunits, SMARCC1, SMARCC2, and SMARCD1/2/3. BAF and PBAF additionally share SMARCB1 and SMARCE1 core subunits. SWI/SNF complexes contain 1 of 2 mutually exclusive ATPase subunits, SMARCA4 and SMARCA2. These isoforms also have distinct sets of isoform-specific subunits which facilitate their occupancy at target gene promoters and enhancers. ARID1A/B and DPF1/2/3 are BAF-specific subunits, ARID2, PBRM1, PHF10, and BRD7 are PBAF-specific subunits, and BICRA/AL and BRD9 are ncBAF-specific subunits^13^.

Mutations in the SWI/SNF complex appear in many adult and pediatric tumors including non-small cell lung cancer, renal clear cell carcinoma, rhabdoid tumors, medulloblastoma, and several types of gynecological cancers. Specific SWI/SNF subunit mutations are often associated with particular cancer types^14–21^. In ovarian cancers, about half of ovarian clear cell carcinomas (OCCC) bear mutations in *ARID1A*^22–24^. High grade serous ovarian carcinomas (HGSOC) bear mutations in a number of SWI/SNF subunits (10% *SMARCA4*, 4% *SMARCA2*, 4% *ARID2*, 2% *SMARCB1*, 2% *ARID1B* and 1% *ARID1A*)^25^. In other gynecological malignancies, *ARID1A* is mutated in ~30% of endometrial carcinomas^22^ and *SMARCA4* is recurrently mutated in undifferentiated uterine sarcomas^26,27^.

Additionally, we and others have discovered that nearly all (>98%) small cell carcinoma of the ovary hypercalcemic type (SCCOHT) tumors are driven by mutations in the *SMARCA4* ATPase^28–31^. The mutually exclusive alternative catalytic subunit, *SMARCA2*, is epigenetically silenced in these tumors^30,32^. SCCOHTs have remarkably stable genomes, with *SMARCA4* being the only recurrently mutated gene. SCCOHT tumors are poorly differentiated and clinically aggressive, occurring in young women (average onset age 24, range 14 months - 43 years)^33,34^. Long term survival of all SCCOHT patients is estimated to be 10-20% with the best outcomes associated with early stage diagnosis and high dose chemotherapy in conjunction with autologous stem cell rescue^34^. Currently, no targeted therapies are available for SCCOHT patients. SMARCA4/A2 dual-deficient SCCOHT cells present a unique model to study the biology of SWI/SNF with regard to its non-catalytic functions.

In order to characterize the effects of SMARCA4/A2 dual loss in SCCOHT on SWI/SNF complex composition, protein-protein interactions (PPIs) and their downstream consequences, we used immunoprecipitation and mass spectrometry (IP-MS) employing 6 antibodies to representative SWI/SNF subunits in 3 SCCOHT cell lines compared to a cell line with a wild-type SWI/SNF complex. We detected an intact core SWI/SNF complex, loss of ATPase module subunits, and loss of PBAF isoform specific subunits in all 3 SCCOHT cell lines. Compared to the wild-type SWI/SNF complex, the SCCOHT SWI/SNF complex showed reduced interactions with proteins involved in chromatin organization and histone post-translational modification. The chromatin regulatory PPIs absent from the SCCOHT SWI/SNF complex include ATPase module and PBAF-specific SWI/SNF subunits, the nucleosome remodeling and deacetylase (NuRD) complex, and the histone methyltransferase MLL3/4 complex.

The SCCOHT SWI/SNF complex also showed exclusive interactions with central metabolism proteins and increased interactions with proteins involved in RNA processing compared to the wild-type SWI/SNF complex. We also observed that a subset of SCCOHT-exclusive SWI/SNF interacting proteins may be synthetic lethal targets. Taken together, we have elucidated altered molecular interactions of the residual SCCOHT SWI/SNF complex which provide insight into the functional consequences of SWI/SNF subunit loss and may identify therapeutic vulnerabilities. As SWI/SNF complex members are mutated in 20% of cancers, these molecular interactions may be relevant for many tumor types.

## Materials and Methods

### Cell culture

BIN67 (BIN-67), SCCOHT1 (SCCOHT-1), and COV434 are SCCOHT cell lines bearing mutations in SMARCA4 and lacking SMARCA2 expression^35–37^. SCCOHT1 and BIN67 cells were maintained in Roswell Park Memorial Institute (RPMI) 1640 medium supplemented with 10% Fetal Bovine Serum (FBS) and 1% Penicillin/Streptomycin (PS). COV434 cells were maintained in Dulbecco’s Modified Eagle Medium (DMEM) supplemented with 10% FBS and 1% PS. HAP1 is a haploid chronic myelogenous leukemia cell line derived from KBM-7 bearing a wild-type SWI/SNF complex (Sequence Read Archive accession SRP044390)^38^. HAP1 cells were maintained in Iscove’s Modified Dulbecco’s Medium (IMDM) supplemented with 10% FBS and 1% PS. SMARCA4 expression was induced in COV434 pIND20-BRG1-2.7 cells by exposure to 500 ng/mL doxycycline for 72 hours^39^. All cell lines were cultured at 37°C in a humidified incubator with 5% CO_2_. Cell lines were Short Tandem Repeat (STR) profiled for verification and monitored routinely for mycoplasma contamination. Cells were harvested by scraping into cold 1× Phosphate Buffered Saline (PBS), followed by centrifugation, flash freezing, and storage at −80°C.

### Immunoprecipitation

Cytoplasmic extracts were first isolated after thawing cells on ice by incubation in 400 μL Buffer A (20 mM HEPES pH 7.9, 10 mM KCl, 0.2 mM EDTA pH 8, 1× Halt™ protease inhibitor cocktail) on ice for 10 minutes. After addition of 25 μL of 10% NP-40, extracts were vortexed, and nuclei were pelleted by centrifugation at 21,000 RCF at 4°C for 30 seconds. Nuclei were resuspended in 400 μL Buffer C (20 mM HEPES pH 7.9, 0.4 M NaCl, 10 mM EDTA pH 8, 1 mM EGTA pH 8, 1× Halt™ protease inhibitor cocktail) and incubated at 4°C for 15 minutes using end-over-end rotation. Insoluble fraction was separated by centrifugation at 21,000 RCF at 4°C for 5 minutes. Soluble nuclear protein extracts were collected and protein concentration was measured using a BCA microplate assay. An input of 500 μg of nuclear protein was used for each IP and the total volume was adjusted to 400 μL. Either protein A or G conjugated sepharose beads, used with rabbit and mouse antibodies, respectively, were washed 3 times in Buffer C, and 30 μL of a 50% slurry were added to appropriate IP samples. A titrated amount of each antibody used was added (see Table 1) to a protein/bead mixture and incubated at 4°C overnight using end-over-end rotation. Sepharose beads were pelleted by centrifugation at 6,000 RCF at 4°C for 1 minute and IP samples were washed 5 times with 500 μL Buffer C.

**Table 1.** Antibodies used for immunoprecipitation in this study.

| Antibody target | Host | Source | Catalogue # | Volume per 500 μg IP |
| --- | --- | --- | --- | --- |
| BAF57/SMARCE1 | Rabbit | Bethyl | A300-810A | 3 μL |
| SMARCB1/SNF5 | Rabbit | Bethyl | A301-087A | 2 μL |
| ARID2 | Rabbit | Cell Signaling | 82342 | 2 μL |
| BAF155/SMARCC1 | Rabbit | Cell Signaling | 11956 | 4 μL |
| ARID1A | Mouse | Millipore | 04-080 | 2 μL |
| BRG1/SMARCA4 | Rabbit | AbCam | ab215998 | 1.5 μL |
| IgG isotype | Mouse | Santa Cruz | sc-2025 | 0.5 μL |
| IgG isotype | Rabbit | Non-commercial | Polysera | 0.5 μL |

### SDS-PAGE and Trypsin Digestion

Proteins were eluted from dried beads by reconstituting in Laemmli loading buffer containing 2-Mercaptoethanol then incubating for 5 minutes at 95°C. Samples were separated by SDS-PAGE using 4-20% Tris-Glycine gels. Proteins were visualized by Coomassie staining, each sample was divided into 4 equal fractions, and each fraction was cut into 2 mm^3^ cubes. Overnight in-gel trypsin digestion was performed following previously published methods^40^. Prior to analysis, peptides were desalted following a modified C18 StageTip protocol^41^. Peptides were resuspended in 2% acetonitrile, 0.1% trifluoroacetic acid (TFA) and loaded onto a self-packed C18 (3M, Maplewood, MN) StageTip. Peptides were washed with 2% acetonitrile, 0.1% TFA, eluted in 60% acetonitrile 0.1% TFA, dried in a vacuum concentrator, and stored at −80°C.

### Isobaric labeling based nuclear proteomics of SCCOHT cell lines

Nuclei were isolated from HAP1, BIN67, COV434 and SCCOHT1 cell lines by resuspending cells in 400 μl buffer (20 mM HEPES pH 7.9, 10 mM KCl, 0.2 mM EDTA pH 8, 1× Halt™ protease inhibitors) and incubated for 10 minutes on ice. After addition of 25 μL of 10% NP-40, extracts were vortexed, and nuclei were pelleted by centrifugation at 21,000 RCF at 4°C for 30 seconds. Resulting nuclei were solubilized in 8M urea lysis buffer with protease inhibitors and sonicated. Protein concentration was determined by BCA assay on clarified nuclear lysates. 200 μg of nuclear protein extract per cell line was reduced by TCEP (10 mM final concentration) and alkylated by iodoacetamide (40 mM final concentration). Samples were first digested overnight by trypsin (1:50 enzyme to substrate ratio) followed by a second trypsin digestion for 1 hour (1:50 enzyme to substrate ratio). Tryptic peptides were desalted using Sep-Pak C18 columns (Waters Inc., Milford, MA) and peptide yield was determined using a BCA assay. Equal amounts of tryptic peptides (90 μg each) were labeled with distinct Tandem Mass Tag (TMT) reagents as per manufacturer’s recommendation (ThermoFisher Scientific). Samples were pooled and desalted following determination of labeling efficiency of >99%. Pooled, TMT-labeled samples were depleted of phosphopeptides using sequential TiO_2_ and Immobilized Metal Affinity Chromatography (IMAC) enrichment strategy and the flow through containing unmodified peptides was subjected to high pH offline fractionation. The resulting 96 fractions were concatenated into 24 fractions for LC-MS/MS analysis^42^.

### Liquid Chromatography and Mass Spectrometry

Peptides were reconstituted in 2% acetonitrile, 0.1% formic acid containing 25 fmol of Pierce Peptide Retention Time Calibration (PRTC) mixture. Data were acquired on an Orbitrap Fusion Lumos (ThermoFisher Scientific, Waltham, MA) interfaced with an Ultimate 3000 UHPLC system (ThermoFisher Scientific) running binary solvent system A (Water, 0.1% formic acid) and B (Acetonitrile, 0.1% formic acid). IP-MS samples (5 μL injection volume) were loaded directly (18.3 minutes loading time) on an analytical column (PepMap RSLC C18, 75 μm ID x 15 cm, 3 μm particle size, 100 Å pore size) kept at 45°C and eluted at a flow rate of 300 nL/minute using the following 60 minutes method: 2% to 19% solvent B in 42 minutes, 19% to 45% B in 6 minutes, 45% to 90% B in 0.5 minute, plateau at 90% B for 1 minute, return to initial conditions in 0.5 minute and re-equilibration for 10 minutes. Data-dependent acquisition was performed in Top Speed mode with a 3 second duty cycle and the following parameters: spray voltage of 1900V, ion transfer tube temperature of 275°C, survey scan in the Orbitrap at a resolution of 120K at 200 m/z, scan range of 400-1500 m/z, AGC target of 4E5 and maximum ion injection time of 50 milliseconds. Every parent scan was followed by a daughter scan using High Energy Collision (HCD) dissociation of top abundant peaks and detection in the iontrap with the following settings: quadrupole isolation mode enabled, isolation window at 1.6 m/z, AGC target of 5E3 with maximum ion injection time of 35 milliseconds and HCD collision energy of 35%. Dynamic exclusion was set to 60 seconds.

Nuclear proteomics samples (5 μL injection volume, ~1 μg peptides per fraction) were loaded directly (18.3 minutes loading time) on an analytical column (PepMap RSLC C18, 75 μm ID x 25 cm, 2 μm particle size, 100 Å pore size) kept at 45°C and eluted at a flow rate of 300 nL/minute using the following 120 minutes method: 2% to 19% solvent B in 80 minutes, 19% to 30% B in 20 minutes, 30% to 98% B in 5 minutes, plateau at 98% B for 2 minutes, return to initial conditions in 1 minute and re-equilibration for 12 minutes. Data-dependent acquisition was carried out using SPS-MS3 workflow in Top Speed mode with a 3 second duty cycle and the following parameters: spray voltage of 2300 V, ion transfer tube temperature of 275°C, survey scan in the Orbitrap at a resolution of 120K at 200 m/z, scan range of 375-1500 m/z, AGC target of 4E5 and maximum ion injection time of 50 milliseconds. Every parent scan was followed by a daughter scan using Collision Induced Dissociation (CID) of top abundant peaks and detection in the iontrap with the following settings: quadrupole isolation mode enabled, isolation window at 0.7 m/z, AGC target of 1E4 with maximum ion injection time of 50 milliseconds and Normalized Collision Energy (NCE) of 35%. Up to 10 MS2 fragment ions selected in the Synchronous Precursor Selection (SPS) node were subjected to MS3 scan. MS3 scan was performed in the Orbitrap at 50,000 resolution over a mass range of 100-500 m/z on MS3 precursors fragmented by HCD (NCE of 65%, AGC target of 1E5 and maximum injection time of 105 milliseconds). Dynamic exclusion was set to 60 seconds.

### Protein Identification

Mass spectra were aligned within each cell line and IP, followed by peak picking using Progenesis QI for proteomics v4.1.6 with default parameters for automated processing. Preprocessed spectra were then searched against a *Homo sapiens* database (UniprotKB/Swissprot, 2017) in Mascot v2.6.0. Oxidation (M) and acetylation (N-terminus) were specified as variable modifications and carbamidomethyl (C) was defined as a fixed modification. MS/MS tolerance was set to 0.6 Da. Tryptic peptides allowed for up to 2 missed cleavages, peptide charge states ranging between +2 and +4, and a peptide tolerance of 10 ppm. Matched spectra were imported into Progenesis for peptide assignment and filtering (Mascot score ≥21). Analyzed gel fractions were recombined and normalized in Progenesis. Normalized abundances of identified proteins were used to score *bona fide* protein-protein interactions (SAINTexpress)^43^ that were enriched at least 2-fold above IgG controls. As per SAINTexpress recommendation, we used a threshold of an average probability of interaction (AvgP) of ≥0.7. Fold change for SWI/SNF complex members in the SCCOHT cell lines was calculated by first taking the mean normalized SAINTexpress abundance for each antibody, then the median across antibodies for a single cell line, and dividing by the same value in HAP1.

For deep expression nuclear proteomics, raw spectra were processed in Proteome Discoverer v2.2 and searched against a *Homo sapiens* protein database (SwissProt/UniprotKB, 2017) in Mascot v2.6.0. Mascot search parameters were the same as above, with quantification performed on reporter ion abundance for isobaric labeling. Identified proteins were filtered for presence in >50% of samples and normalized using the SL and TMM normalization functions from the limma package (v3.38.3) in R v3.5.1.

### Western blotting

Western blotting was performed using 30 μg of protein extracts from BIN67, SCCOHT1, COV434, HAP1, and COV434 pIND20 BRG1-2.7 cells extracted in either nuclear protein extraction buffers described above or whole cell lysis buffer (0.05 M Tris-HCl pH 6.8, 2% SDS, 6% β-mercaptoethanol) and boiled for 5 minutes. Proteins were separated by 3-8% Tris-Acetate SDS-PAGE and transferred to nitrocellulose membranes. Transfer efficiency was monitored by either REVERT total protein stain or Ponceau stain and was used for normalization. SWI/SNF subunits were detected using primary antibodies to PBRM1 (Bethyl A301-591A, 1:2000), SMARCA4 (AbCam ab110641, 1:2000), SMARCC1 (Cell Signaling 11956, 1:1000), ARID1A (Millipore 04-080, 1:1000), SMARCA2 (Cell Signaling 6889, 1:1000), SMARCE1 (AbCam ab131328, 1:1000), SMARCB1 (Bethyl A301-087A, 1:1000), and ARID2 (Thermo PA5-35857, 1:5000) in either 5% non-fat dry milk or LiCor Odyssey blocking buffer. Protein detection was performed either using horseradish peroxidase-conjugated secondary antibodies and chemiluminescence developed on film or near-infrared fluorescent secondary antibodies developed on a LiCor CLx instrument.

### DNA sequencing and variant calling

Genomic DNA isolation and library preparation was carried out as described previously^28^, using DNeasy Blood & Tissue kit (Qiagen). The precapture libraries were prepared using a combination of On-Bead Whole Genome Sequencing (Kapa) kit and SureSelect XT2 (Agilent) adapters. SureSelect XT2 was further used for capture and clean-up. Paired end sequencing was performed on the Illumina HiSeq 2000. Fastq reads were aligned to the hg19 human genome using BWA-MEM version 0.7.8, followed by sorting and indexing the bam files (Picard version 1.128), duplicate marking, indel realignment and recalibration using GATK version 3.3.0. Variants were called using Haplotype Caller and VCF annotation was done using SNPEff version 3.5h. Protein coding variants that passed the HaplotypeCaller quality filter, classified as “rare” in 1000 Genomes Project and ExAC databases as well as annotated as “High” or “Moderate” impact in SNPEff were considered. Lollipop plots were generated using the trackViewer Bioconductor package in R (version 1.14.1) and Adobe Illustrator.

### Gene Ontology (GO) Biological Processes Analysis

Enriched biological processes were determined using ClueGO v2.3.2 against the *Homo sapiens* database for GO biological processes v06.11.2016_23h27 using experimental evidence only^44^. GO term fusion was applied and only terms with a p-value ≤0.05, based on a 2-sided hypergeometric test with Bonferroni correction, were reported. Kappa score (>0.4) based GO term grouping was utilized to define overarching functional groups across the data. All other parameters were left at default ClueGO settings. Enriched biological processes were merged across the 4 cell lines and ranked in increasing order by term p-value (lowest value among all cell lines). High confidence interactors from all cell lines were then assigned a single GO term from the enriched biological processes based on GO term hierarchy and ClueGO term grouping, and terms without assigned interactors were removed from results (Supplemental Tables 1 & 3).

### STRING Functional Protein Association Networks Analysis

Protein-protein interaction network and enrichment analysis was carried out using stringApp (1.4.2) within Cytoscape (3.7.1)^45^. Uniprot identifiers of proteins uniquely identified in SCCOHT cell lines, HAP1 control cell line or shared between all the 4 cell lines were loaded as separate lists into stringApp and queried against StringDB for known interactions. Resulting interactions were filtered with a minimum confidence (score) cutoff of 0.7 and maximum additional interactors was set to 0. Enrichment analysis was performed using default stringApp settings in Cytoscape.

### CRISPR analysis

Effect on SCCOHT cell growth by CRISPR knock-out of SCCOHT exclusive SWI/SNF interacting proteins was evaluated using Avana CRISPR data available in DepMap (DepMap, Broad (2019): <u>DepMap Achilles 19Q1 Public.</u> figshare. Fileset. doi:10.6084/m9.figshare.7655150)^46^. The CERES score for each of these genes in COV434 and BIN67 cell lines were compared to the average effect of each gene knock-out across all 625 cell lines in the database using a Z score calculated as Z=(x − μ)/σ where x is the CERES score for the gene of interest in either COV434 or BIN67, μ is the average CERES score for the gene of interest across all cell lines in the database, and σ is the standard deviation of the CERES score for the gene of interest across all cell lines in the database.

### Cell viability after SMARCB1 siRNA transfection

Cells were reversely transfected with siRNAs to knock down SMARCB1 (#1: 5’-GCAACGAUGAGAAGUACA -3’ and #2: 5’-GGCCGAGACAAGAAGAGA-3’) and a negative non-targeting control (Integrated DNA technology, 51-01-14) using Lipofectamine RNAiMAX at a final concentration of 5 nM. Knock-down efficiency was evaluated by western blotting 48 hours following transfection. The viability of BIN67 cells was reduced too drastically to provide sufficient material for western blotting. Five days after transfection, cells were fixed with 10% methanol, 10% acetic acid, and stained with 0.5% crystal violet in methanol. The absorbed dye was solubilized in 10% acetic acid and the absorbance was measured at 595 nm in a microplate spectrophotometer. Cell survival was determined by normalizing absorbance values to the non-targeting control treated cells^47^.

## Results

### The SWI/SNF complex in SCCOHT cell lines shows reduced ATPase module and PBAF subunit interactions

Aside from dual loss of SMARCA4 and SMARCA2, knowledge about the SWI/SNF complex composition, PPIs, and downstream consequences of these alterations in SCCOHT cells is limited. As a critical step toward defining the molecular biology of SCCOHT, we characterized the residual SWI/SNF complex in SCCOHT cells by performing IP-MS in 3 SCCOHT cell lines compared to the HAP1 cell line bearing a wild-type SWI/SNF complex. We used 6 antibodies to representative SWI/SNF subunits (ARID1A, ARID2, SMARCA4, SMARCB1, SMARCC1, and SMARCE1) and analyzed complex composition and PPIs by probabilistic scoring using SAINTexpress^43^. Based on SAINTexpress guidelines, we used an interaction confidence threshold of a 2-fold enrichment over IgG and an average probability of interaction (AvgP) ≥ 0.7 with at least 1 antibody. We detected all previously identified mammalian SWI/SNF subunits^48–50^ in the SWI/SNF wild-type cell line HAP1, 26 of which passed our interaction threshold (Figure 1). Core SWI/SNF subunit (SMARCC1, SMARCC2, SMARCD1/2, SMARCB1, and SMARCE1)^13^ interactions were detected above the AvgP threshold in all 3 SCCOHT cell lines, albeit at a lower protein abundance than HAP1 (Figure 1A, processed data in Supplemental Table 1, fold change in Supplemental Table 2).

**Figure 1.**
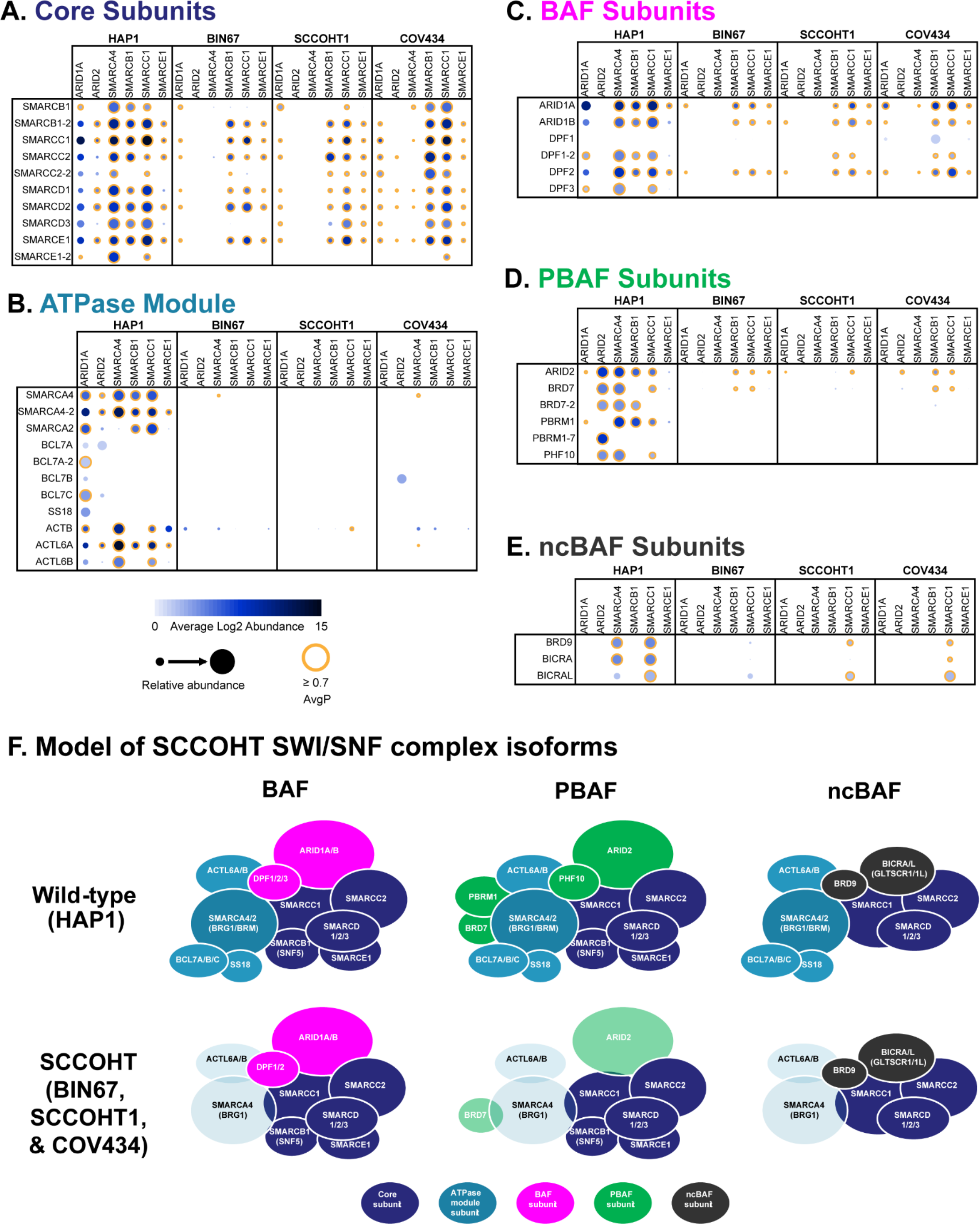
Characterization of SWI/SNF complex integrity in SCCOHT cell lines. **A-E.** Dot plot visualization of IP-MS data from antibodies to 6 different SWI/SNF complex members serving as baits (columns). The selected table of prey (rows) include all previously characterized SWI/SNF complex members, which are grouped as core subunits (A), ATPase module subunits (B), BAF-specific subunits (C), PBAF-specific subunits (D), and ncBAF-specific subunits (E). Bait-prey interaction probabilities were scored by SAINTexpress (AvgP) on a scale of 0-1 and visualized using ProHits-Viz from 3 replicates of each IP from HAP1 (wild-type SWI/SNF), BIN67, SCCOHT1, and COV434 nuclear extracts. Proteins with AvgP ≥0.7 were considered high confidence interactors (orange ring). Log2 abundance is represented by dot color and size, with the size being normalized for each prey (row), across all IPs. **F.** Model of SCCOHT SWI/SNF complex isoforms based on data in A-E. The absence of a protein in SCCOHT IP samples is diagrammed by the absence of the corresponding protein bubble in the cartoon. Reduced protein abundance in SCCOHT IPs compared to HAP1 is diagrammed as a transparent bubble in the cartoon. Core subunits are colored royal blue, ATPase module subunits are colored teal, BAF-specific subunits are colored magenta, PBAF-specific subunits are colored green, and ncBAF-specific subunits are colored dark gray.

Overall, ATPase module subunits are either absent from or reduced in the SCCOHT SWI/SNF complex. Consistent with previous reports that *SMARCA2* is epigenetically silenced in SCCOHT tumors, we did not detect any SMARCA2-specific peptides in any of the 3 SCCOHT cell lines (Figure 1B). Despite the presence of inactivating mutations (nonsense and frameshift mutations in SCCOHT1, 2 splice site mutations in BIN67, and missense and splice site mutations in COV434), we detected SMARCA4-specific peptides in all 3 SCCOHT cell lines at a low abundance (Supplemental Tables 1-2). In BIN67 and SCCOHT1, no other SWI/SNF subunits were observed to interact with SMARCA4 above the confidence threshold (Figure 1B). BCL7A/B/C and SS18 proteins are not detected above the AvgP threshold for interaction in the SCCOHT SWI/SNF complexes. Additionally, the association of SWI/SNF with actin is reduced in SCCOHT cells (Figure 1B, Supplemental Table 2).

Across all 3 SCCOHT cell lines, we observe a decrease in association of PBAF-specific subunits compared to BAF- and ncBAF-specific subunits in the SCCOHT SWI/SNF complex compared to the wild type. The SCCOHT BAF contains a mixture of ARID1A/B and DPF1/2, but lacks DPF3 (Figure 1C). The SCCOHT PBAF has reduced levels of ARID2 and BRD7 and lacks PBRM1 and PHF10 (Figure 1D, Supplemental Table 2). We previously detected PBRM1 RNA expression in SCCOHT cell lines, patient derived xenografts, and tumors above that observed in normal ovaries^30^. However, PBRM1 protein expression was very low in all 3 SCCOHT cell lines by western blotting (Figure 2D). PBRM1 protein levels were increased by either treatment with the proteasome inhibitor, MG132, or by re-expression of SMARCA4, suggesting that PBRM1 protein is not stable in the absence of SMARCA4 (Figure 3). Thus, SCCOHT cells showed a stronger reduction in association with PBAF-specific subunits than BAF-specific subunits. We also observed ncBAF subunits in all 3 SCCOHT cell lines (Figure 1E, Supplemental Table 2). Consistent with previous reports that ncBAF does not include SMARCB1 or SMARCE1, the ncBAF specific subunits BRD9 and BICRA/AL were only detected with the SMARCC1 antibody. The association of SWI/SNF subunits with the residual complex in SCCOHT mirrored the nuclear abundance of each subunit (Figure 2). Based on these data, a model of SWI/SNF composition in SCCOHT is depicted in Figure 1F.

**Figure 2.**
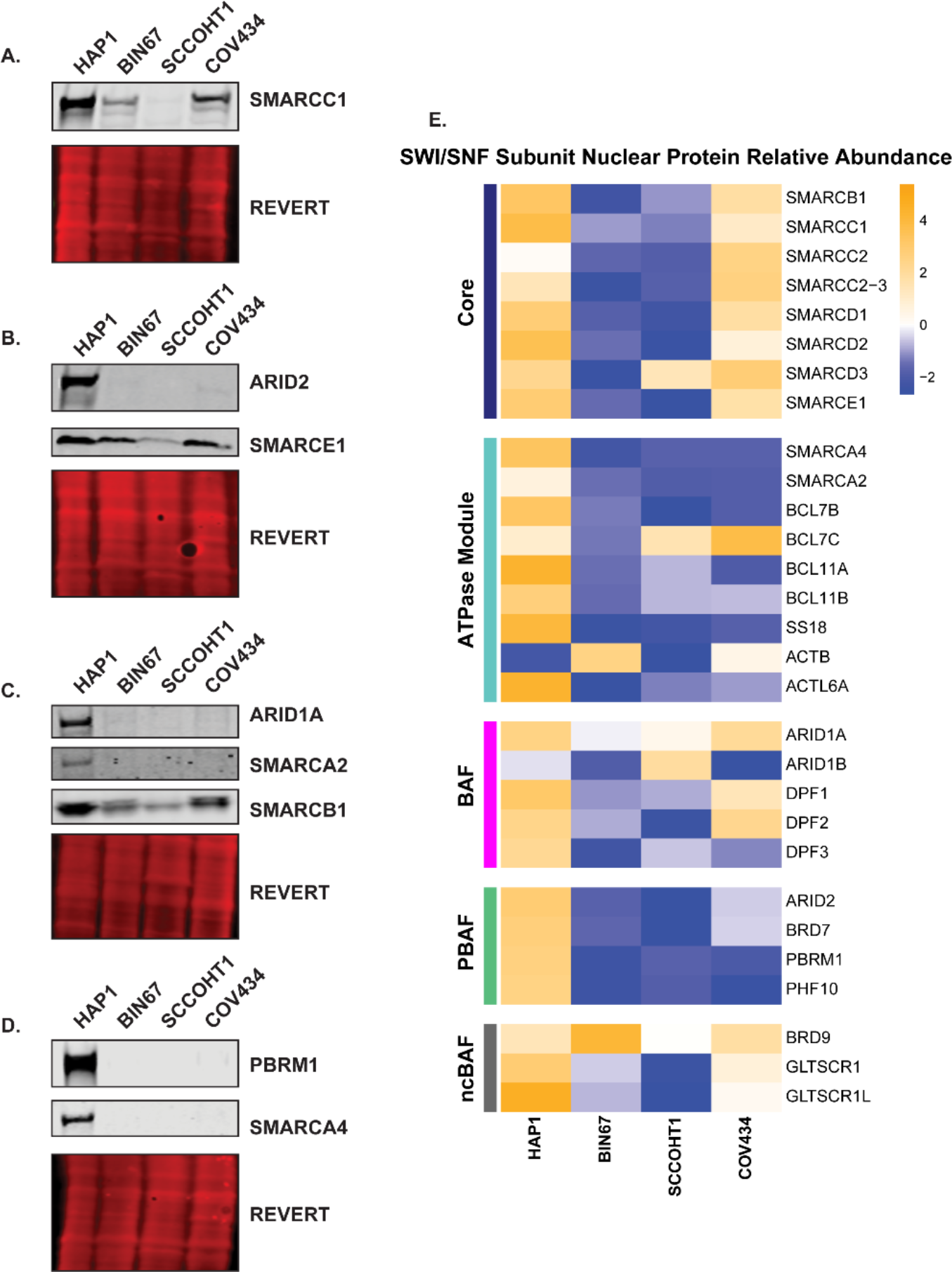
SWI/SNF subunit protein abundance in SCCOHT nuclear extracts. **A.** Representative western blot analysis of SMARCC1 protein levels in HAP1, BIN67, SCCOHT1, and COV434 cell lines compared to total protein transferred as indicated by REVERT stain. **B.** Representative western blot analysis of ARID2 and SMARCE1 protein levels as in (A). **C.** Representative western blot analysis of ARID1A, SMARCA2, and SMARCB1 protein levels as in (A). **D.** Representative western blot analysis of PBRM1 and SMARCA4 protein levels as in (A). **E.** Relative nuclear protein abundance of SWI/SNF subunits measured by LC-MS/MS of trypsin digested, TMT labeled total nuclear protein extracts from HAP1, BIN67, SCCOHT1, and COV434 cells. Heatmap values represent z-scores calculated across the 4 cell lines.

**Figure 3.**
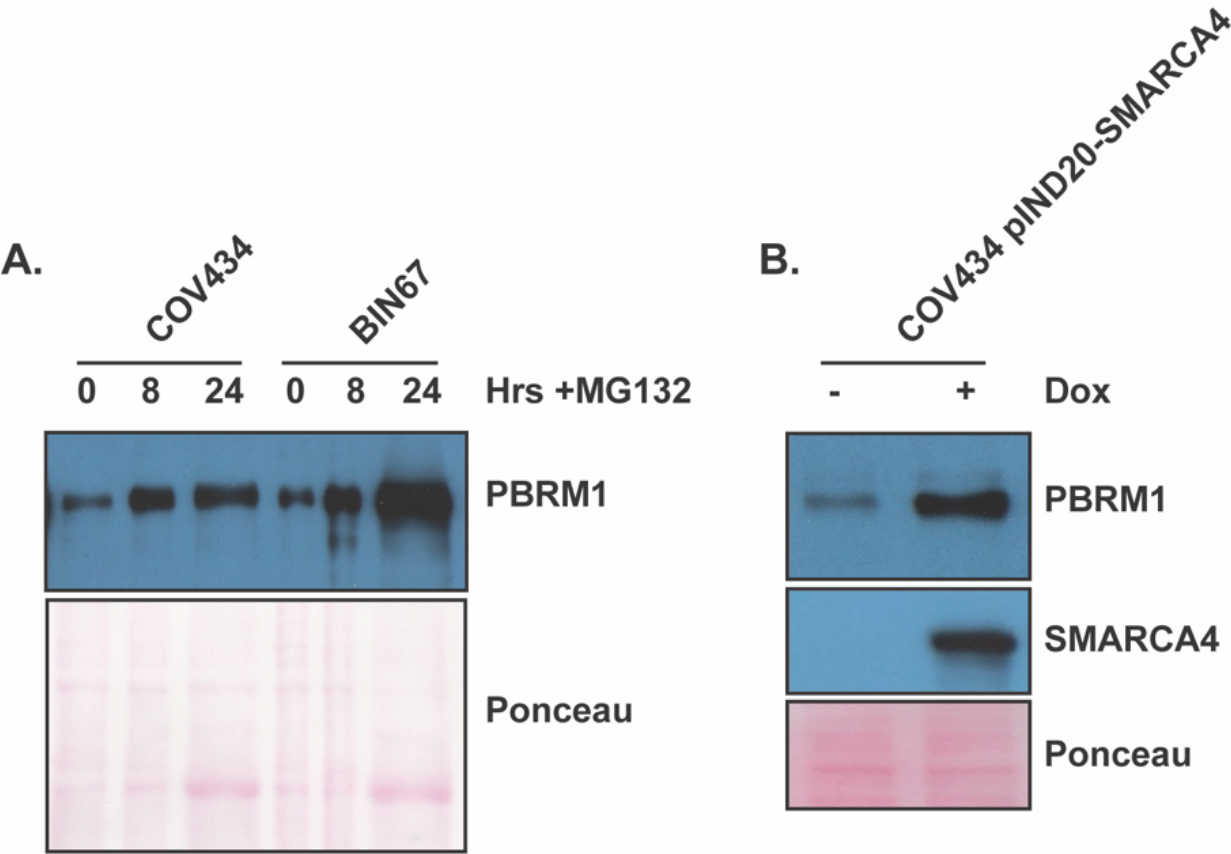
PBRM1 protein is degraded in the absence of SMARCA4 expression in SCCOHT cells. **A.** Western blot analysis of long exposure PBRM1 levels in BIN67 and COV434 whole cell extracts from cells treated with 1 μM MG132 for 0, 8, and 24 hours compared to total protein transferred as indicated by Ponceau stain. **B.** Representative western blot analysis of PBRM1 and SMARCA4 levels as in (A) of COV434 cells re-expressing SMARCA4 under control of the doxycycline inducible promoter, pIND20 (COV434 pIND20-BRG1-2.7).

### Low abundance peptides unique to SMARCA4 are expressed in SCCOHT cell lines

As *SMARCA4* is mutated in the SCCOHT cell lines and its expression was undetected by western blotting, we analyzed LC-MS/MS peptide coverage with respect to where the mutations lie within the protein sequence. The HAP1 cell line is wild-type for all SWI/SNF complex subunits based on prior exome sequencing (Sequence Read Archive accession SRP044390). SMARCA4 peptides are detected throughout the entire protein sequence from HAP1 cells with all 6 antibodies used in this study (Figure 4). The SCCOHT1 cell line contains a nonsense and a frameshift mutation, both occurring in the helicase domain, presumably disrupting ATPase and nucleosome sliding activities^29^. Two SMARCA4 peptides were detected in SMARCA4 IPs from SCCOHT1 cells, both mapping to sites N-terminal to the mutation sites (Figure 4). The BIN67 cell line contains 2 splice site mutations within the Snf2_N domain that would likely disrupt catalytic activity^29^. SMARCA4-specific peptides were detected in the BIN67 SMARCA4 IP that lie N-terminal to the mutations, but no peptides were detected in regions spanning the exons presumed affected by the splice site mutations (Figure 4). The COV434 cell line bears a missense mutation between the QLQ and HSA domains and a splice site mutation in the SnAC domain (A. Karnezis, unpublished). SMARCA4-specific peptides were sparsely detected throughout the sequence of the protein in SMARCA4, ARID1A, SMARCC1, and SMARCB1 IPs from COV434 cells that lie N-terminal to the splice site mutation (Figure 4). Thus, in all SCCOHT cell lines, we detected residual SMARCA4 peptides. In the COV434 cell line, the SMARCA4 antibody immunoprecipitated SMARCA4 and core SWI/SNF complex members, SMARCB1, SMARCC1, SMARCD1/2, and SMARCE1, indicating that SMARCA4 is associated with the SWI/SNF complex in COV434 cells.

**Figure 4.**
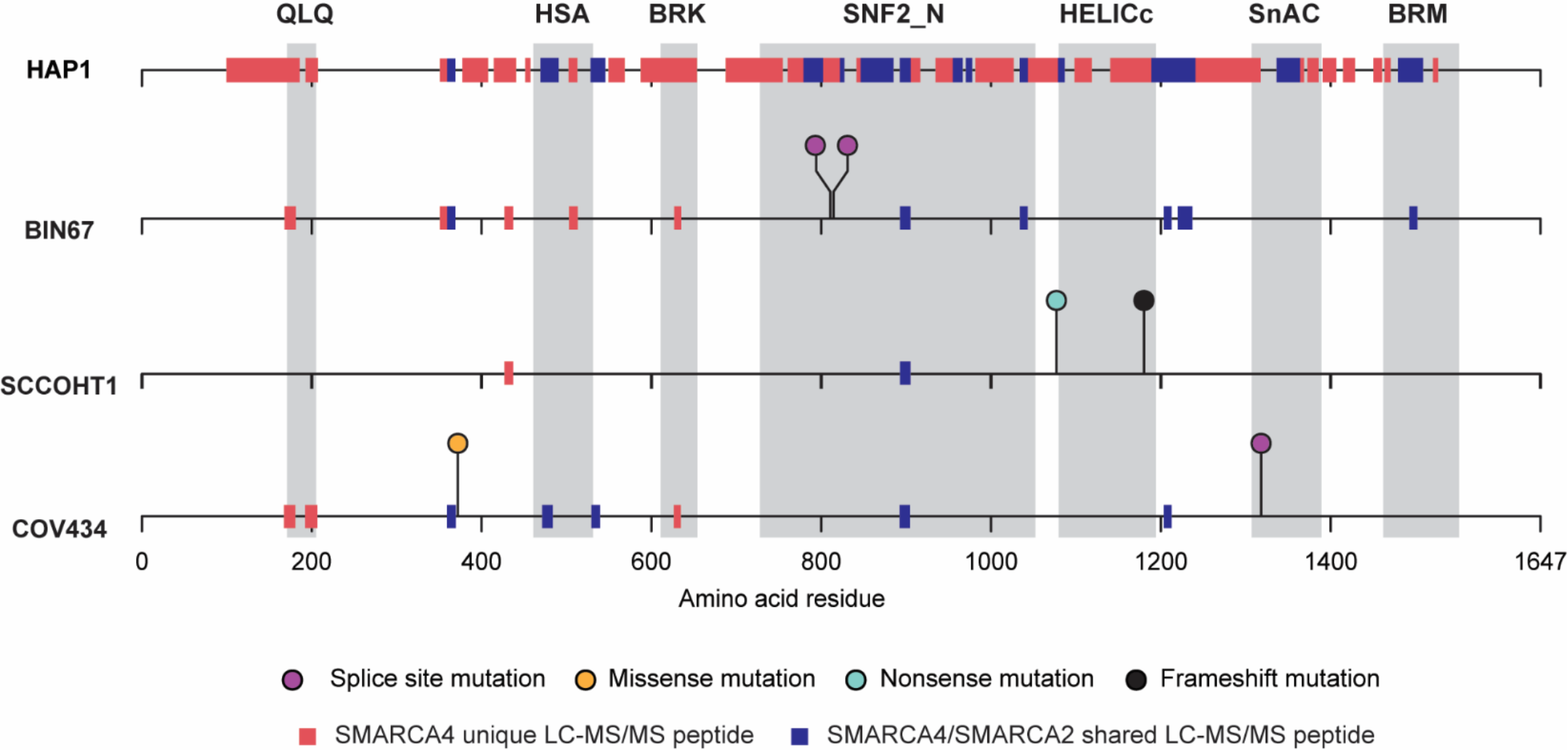
SMARCA4 peptides detected in SCCOHT cell lines. Red boxes on the ruler represent peptides unique to SMARCA4 and blue boxes represent peptides shared by SMARCA4/A2 identified by LC-MS/MS of proteins immunoprecipitated using antibodies against ARID1A, ARID2, SMARCA4, SMARCB1, SMARCC1, and/or SMARCE1 for HAP1, BIN67, SCCOHT1, and COV434. Lollipops above a ruler represent *SMARCA4* genomic alterations identified by WES and RNAseq superimposed onto Pfam SMARCA4 protein domains (gray background).

### Residual SWI/SNF complex is essential for SCCOHT cell viability

To determine whether the residual SWI/SNF complexes observed in SCCOHT are important for cell viability, we assayed cell growth after siRNA knock-down of a core SWI/SNF subunit, SMARCB1. Upon knock-down of SMARCB1 by either of 2 siRNAs, BIN67, COV434, and SCCOHT1 cells showed significantly reduced growth (10-12%, 43-65%, and 13-25% cell growth, respectively) compared to a non-targeting control siRNA (100% cell growth) (Figure 5A & B). These data indicate that even in the context of a residual SWI/SNF complex, loss of SMARCB1 negatively affects growth of SCCOHT cells.

**Figure 5.**
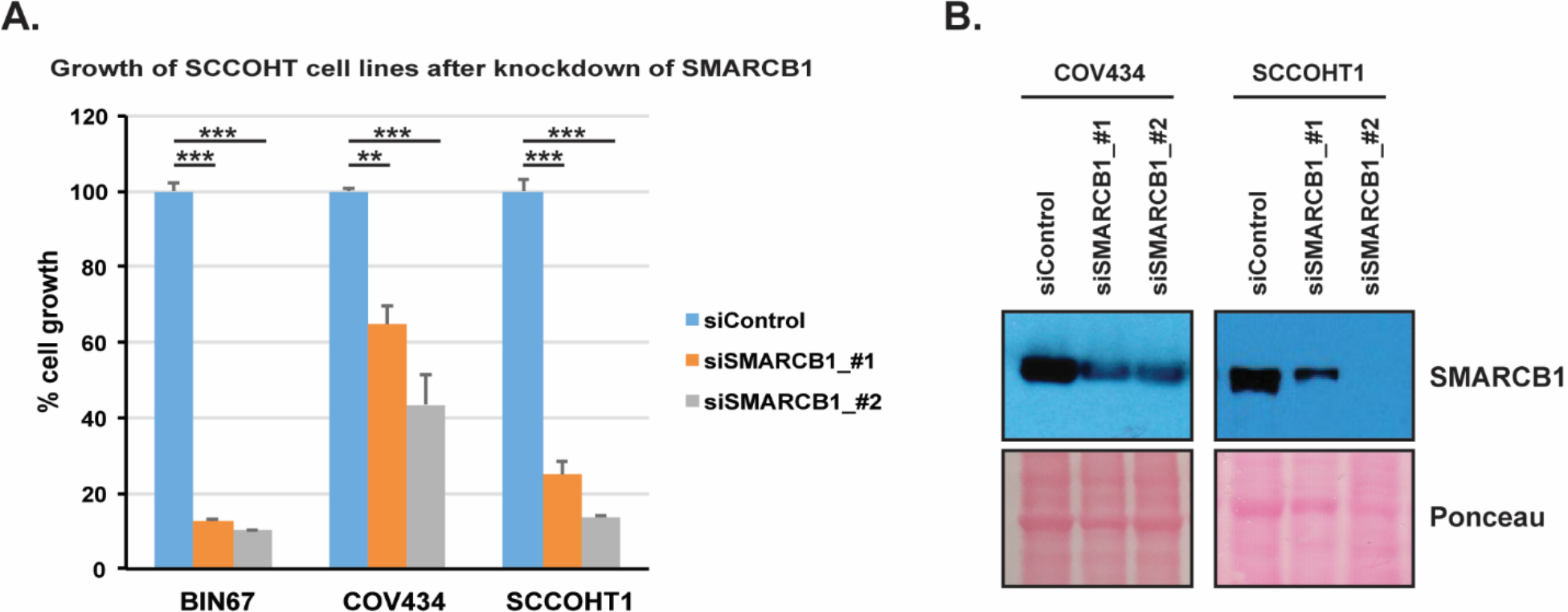
The core SWI/SNF subunit, SMARCB1, is important for SCCOHT cell growth. **A.** Cell growth of BIN67, SCCOHT1, and COV434 cell lines comparing 2 siRNAs knocking down SMARCB1 expression to a control siRNA measured by crystal violet staining 5 days after reverse transfection with the indicated siRNA sequence. Error bars represent the standard deviation of 4 replicates (**, p<0.01; ***, p<0.001). **B.** Representative western blot analysis of SMARCB1 protein levels after knock-down in (A). The viability of BIN67 cells was reduced so drastically that remaining protein levels were insufficient for western blotting.

### The SCCOHT SWI/SNF complex shows increased interactions with RNA processing proteins and decreased interactions with chromatin regulation proteins

Genetic inactivation and protein expression loss of SWI/SNF subunits in cancer has been associated with broad shifts in SWI/SNF interaction networks that can fuel malignant phenotypes^51^. In order to assess potentially altered PPIs due to SMARCA4 loss in SCCOHT, we analyzed SWI/SNF PPI networks combining data from each of the 6 SWI/SNF subunit IPs and applied ClueGO Gene Ontology (GO) analysis to determine significant enrichment for biological processes for each cell line (Figure 6A-D, Supplementary Table 3). Proteins involved in translation, histone post-translational modification, telomere maintenance, viral response, chromatin organization, cell cycle progression, and RNA processing were observed as significantly enriched categories across the 4 cell lines (Figure 6E, GO IDs provided in Supplementary Table 1). The majority of observed SWI/SNF PPIs are noted as “Unclassified” as they were not assigned to GO biological processes that were significantly enriched across all cell lines.

**Figure 6.**
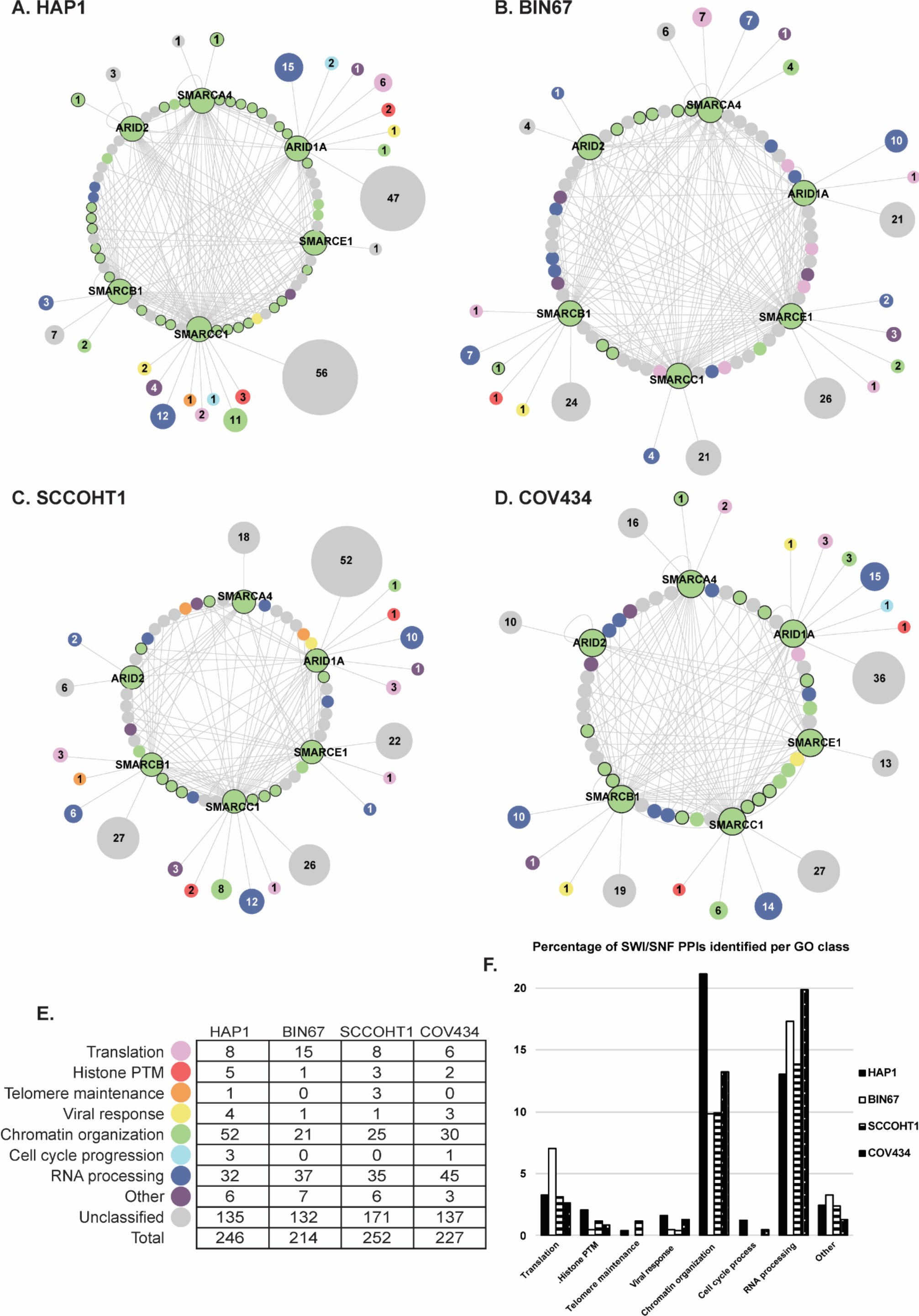
The SCCOHT SWI/SNF complex has more PPIs with RNA processing proteins and less PPIs with chromatin organization proteins. **A-D.** SWI/SNF PPI networks for all interacting proteins (nodes) with an AvgP ≥ 0.7 using antibodies recognizing the ARID1A, ARID2, SMARCA4, SMARCB1, SMARCC1, and SMARCE1 SWI/SNF subunits in the HAP1 (**A**), BIN67 (**B**), SCCOHT1 (**C**), and COV434 (**D**) cell lines. Circles with protein names represent the bait proteins used for immunoprecipitation. Enriched GO terms were ranked by p-value and condensed into overlapping hierarchical groups across all cell lines such that each protein would be assigned to a single GO term. Five GO terms were enriched in at least 1 cell line (kidney development, response to unfolded proteins, substantia nigra development, platelet aggregation and negative regulation of transforming growth factor beta receptor signaling pathway) that were not associated with a parental GO term and were classified as “Other.” GO term colors and numbers of proteins in each category are listed in (E). Nodes in the inner ring indicate proteins that were immunoprecipitated with 2 or more antibodies. Nodes outside the ring represent proteins immunoprecipitated with only 1 antibody which have been collapsed into 1 node for each GO biological process with the size of the node being scaled to the number of proteins belonging to that biological process. Proteins outlined with a gray circle are previously known SWI/SNF complex members. Proteins that were not assigned to a significantly enriched GO term were categorized as unclassified. **E**. Tabulation of the number of SWI/SNF interacting proteins assigned to each enriched GO biological process for the HAP1, BIN67, SCCOHT1, and COV434 cell lines. **F**. Percentage of PPIs in each GO category as in (E), detailed in Supplemental Table 1.

We analyzed differences in the representation of GO biological processes as the percentage of PPIs falling into each process for each cell line. We detected fewer SWI/SNF-interacting proteins involved in histone post-translational modification and chromatin organization in the 3 SCCOHT cell lines compared to HAP1 (Figure 6F). We observed an increase in PPIs with proteins involved in RNA processing in all 3 SCCOHT cell lines compared to HAP1 (Figure 6F). Similarly, we observed that several of the RNA processing proteins were present in at least 2 of 3 SCCOHT cell lines and absent from HAP1 (Figure 7) as described below. We also observed more RNA processing proteins being captured by more than 1 IP antibody in the SCCOHT cell lines than the wild type (Figure 6A-D). The wild-type PBAF complex (immunoprecipitated with ARID2) did not show interactions with RNA processing proteins. However, the SCCOHT PBAF complex interacted with RNA processing proteins in all 3 SCCOHT cell lines (Figure 6A-D).

**Figure 7.**
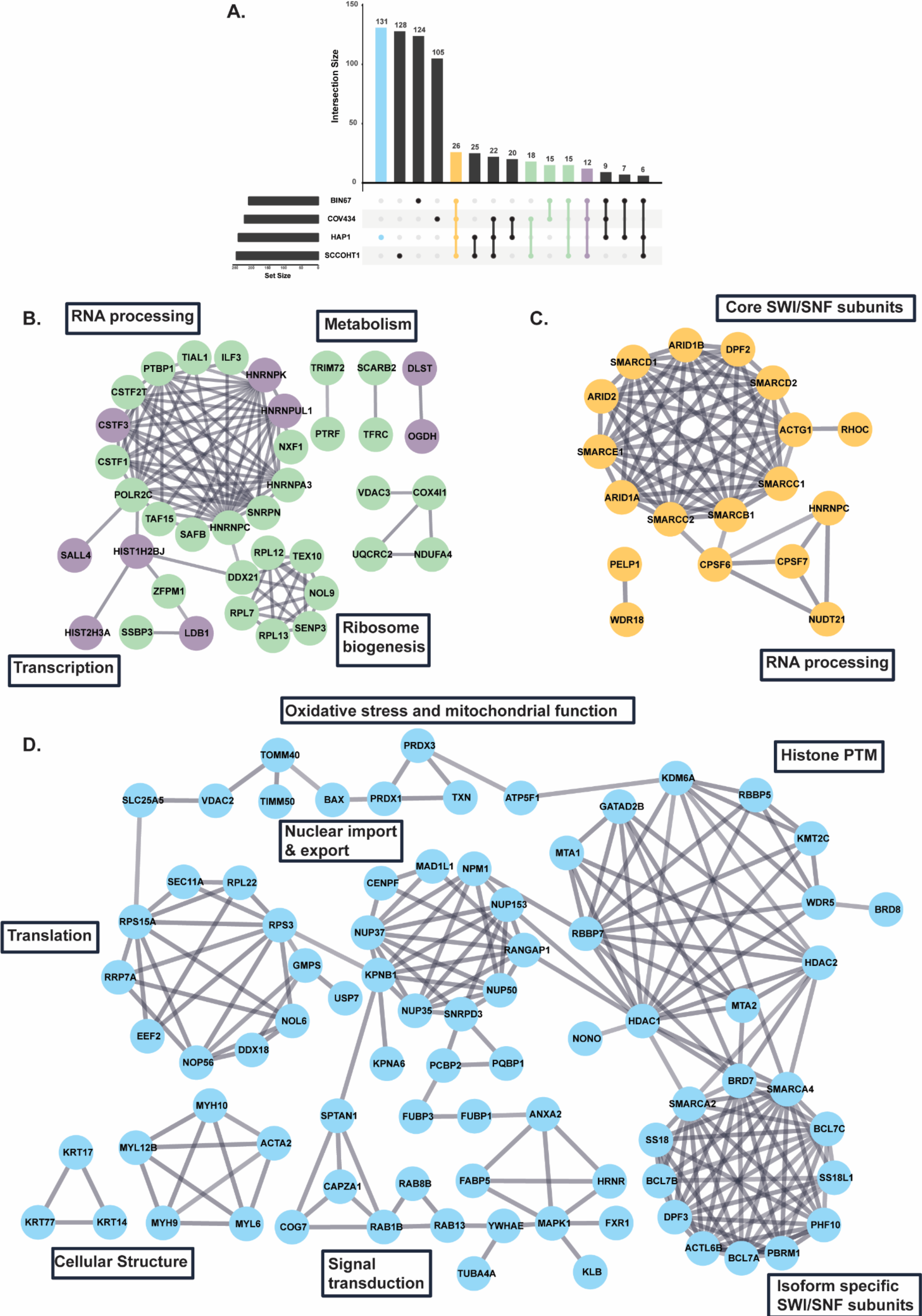
The wild-type and SCCOHT SWI/SNF complexes interact with distinct protein networks. **A**. UpSet plot comparing cumulative overlap of SWI/SNF PPIs (AvgP≥ 0.7) from IP-MS data across 6 SWI/SNF antibodies comparing HAP1, BIN67, SCCOHT1, and COV434. **B**. Interaction network of proteins identified in more than 1 SCCOHT cell line but not observed in HAP1 parental as shown in (A) (sum of 60 proteins) based on previously published literature using StringDB. Proteins interacting with SWI/SNF in 2 or 3 SCCOHT cell lines are colored green or purple, respectively. **C**. Interaction network of SWI/SNF interacting proteins identified in all cell lines (26 proteins, orange) as in (B). **D**. Interaction network of SWI/SNF interacting proteins identified exclusively in HAP1 (131 proteins, blue) as in B-C. For clarity, proteins showing no first-degree interactions in StringDB are not shown (B-D).

To determine what altered interaction networks are unique to SMARCA4-deficient SWI/SNF complex isoforms in SCCOHT cell lines, we compared their PPI networks to the wild-type SWI/SNF complex isoforms expressed in the HAP1 cell line. We identified 664 SWI/SNF-interacting proteins across all cell lines. We observed 214-253 SWI/SNF-interacting proteins in each cell line, with about half of those (105-131) being unique to each cell line. We observed 26 interactions that were shared among all cell lines and 60 interactions that were shared only among 2-3 SCCOHT cell lines (Figure 7A, Supplementary Table 4).

To further explore these relationships, we performed evidence-based functional analysis using interactions previously documented in STRING-DB^52^. PPI networks exclusive to the residual complex and shared in at least 2 of the 3 SCCOHT cell lines were enriched for RNA processing, metabolism, ribosome biogenesis, and transcription (Figure 7B). There were 26 SWI/SNF-interacting proteins that were shared among all cell lines including 12 SWI/SNF complex members, components of the PELP1 complex, and RNA processing factors (Figure 7C). SWI/SNF PPIs found only in the wild-type cell line (131 proteins) (Figure 7D) were associated with oxidative stress and mitochondrial function, histone post-translational modification, translation, nuclear import and export, cellular structure, signal transduction, and isoform-specific SWI/SNF subunits.

The reduction in SWI/SNF-interacting proteins involved in chromatin organization in SCCOHT cells compared to the wild-type cell line can be explained by the absence of several isoform-specific SWI/SNF subunits, including PBAF subunits. Additionally, GO enrichment analysis for cellular components revealed several chromatin regulatory complexes among the SWI/SNF interacting proteins only detected in the wild-type cell line. These include the nucleosome remodeling and deacetylase (NuRD) complex, the MLL3/4/COMPASS complex, the microtubule cytoskeleton, and the nuclear pore basket (Supplementary Table 5).

As described above, the residual SCCOHT SWI/SNF complex showed increased interactions with RNA processing proteins, many of which were exclusive to the SCCOHT SWI/SNF complex. The majority of RNA processing proteins observed were exclusive to 1 cell line (19 to COV434, 15 to BIN67, 9 to SCCOHT1, and 10 to HAP1), 7 proteins were shared across all cell lines, and 14 proteins were observed in 2-3 SCCOHT cell lines (Figure 8). Further comparison of the SWI/SNF-interacting RNA processing proteins between the SCCOHT and wild-type cell lines indicate that these proteins have similar GO biological processes including ribosome biogenesis, splicing, and 3’ end formation factors. Additionally, GO cellular component analyses did not identify any protein complexes or components that were differential between RNA processing proteins interacting with SWI/SNF in wild-type or SCCOHT cells (Supplemental Table 6). Instead, the increased number of interacting RNA processing proteins appear to have the same functions as the RNA processing proteins that were shared with the wild-type SWI/SNF.

**Figure 8.**
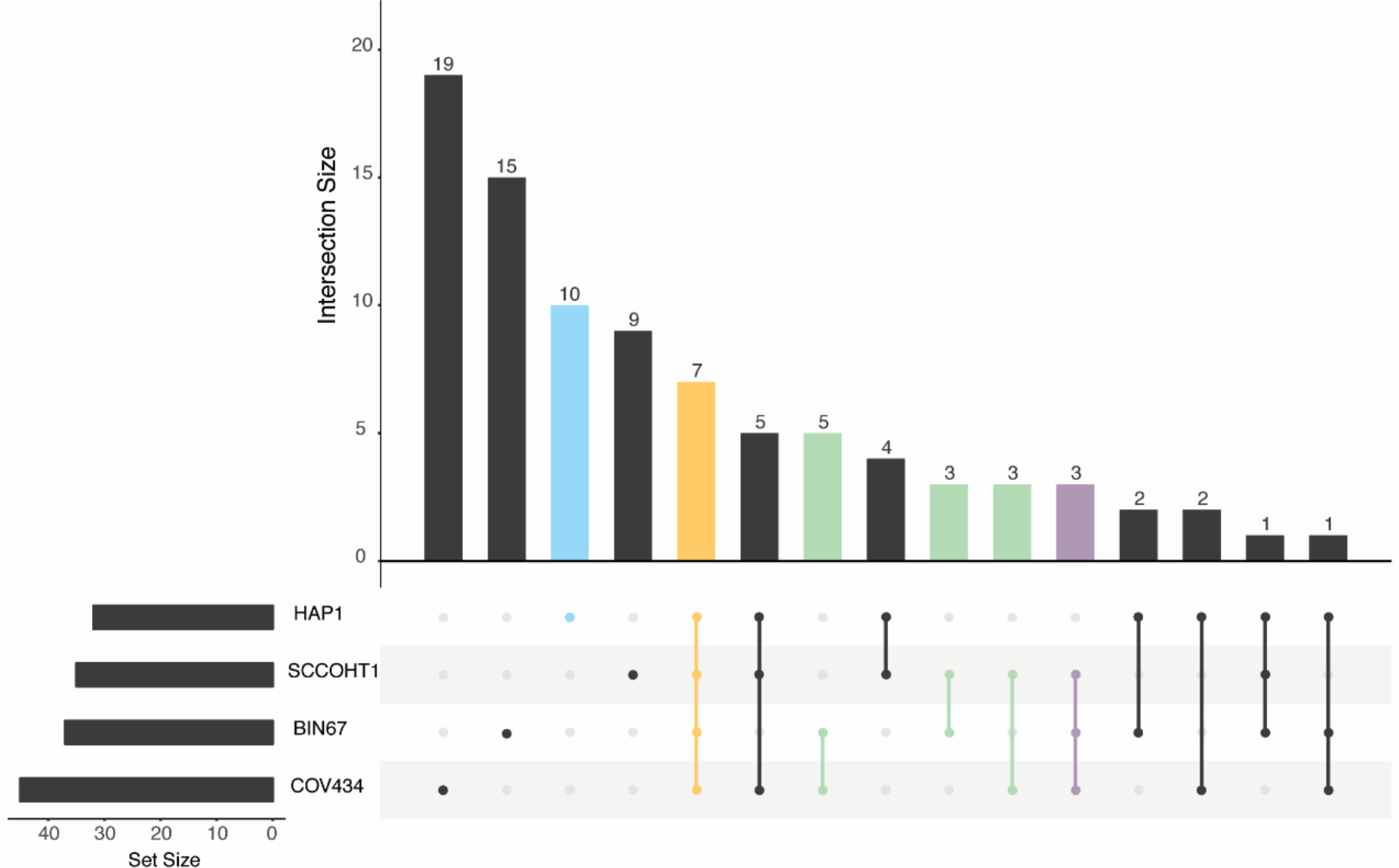
Overlap of SWI/SNF interacting RNA processing proteins. UpSet plot comparing cumulative overlap of SWI/SNF PPIs with RNA processing proteins (AvgP≥ 0.7) from IP-MS data across 6 SWI/SNF antibodies comparing HAP1, BIN67, SCCOHT1, and COV434.

### A subset of proteins interacting exclusively with SCCOHT SWI/SNF isoforms proteins have synthetic lethal dependencies

In order to determine synthetic lethal dependencies of protein interactors exclusive to the residual SWI/SNF in SCCOHT cells, we analyzed how CRISPR knock-out of genes encoding SCCOHT-exclusive SWI/SNF-interacting proteins affected growth of SCCOHT cell lines compared to the average effect across all cell lines in the database using publicly available data^46^. Depletion of 6 genes, highlighted in Figure 9, selectively reduced growth of both BIN67 and COV434. The proteins encoded by those genes include 2 electron transport chain components (UQCRC2 and COX4I1), 2 proteins involved in translation (RPL12 and DDX21), and 2 proteins involved in RNA processing (CSTF3 and PTBP1) (Figure 9). These results may indicate an increased dependency of SCCOHT cells on metabolism, translation, and RNA processing that could guide future therapeutic development.

**Figure 9.**
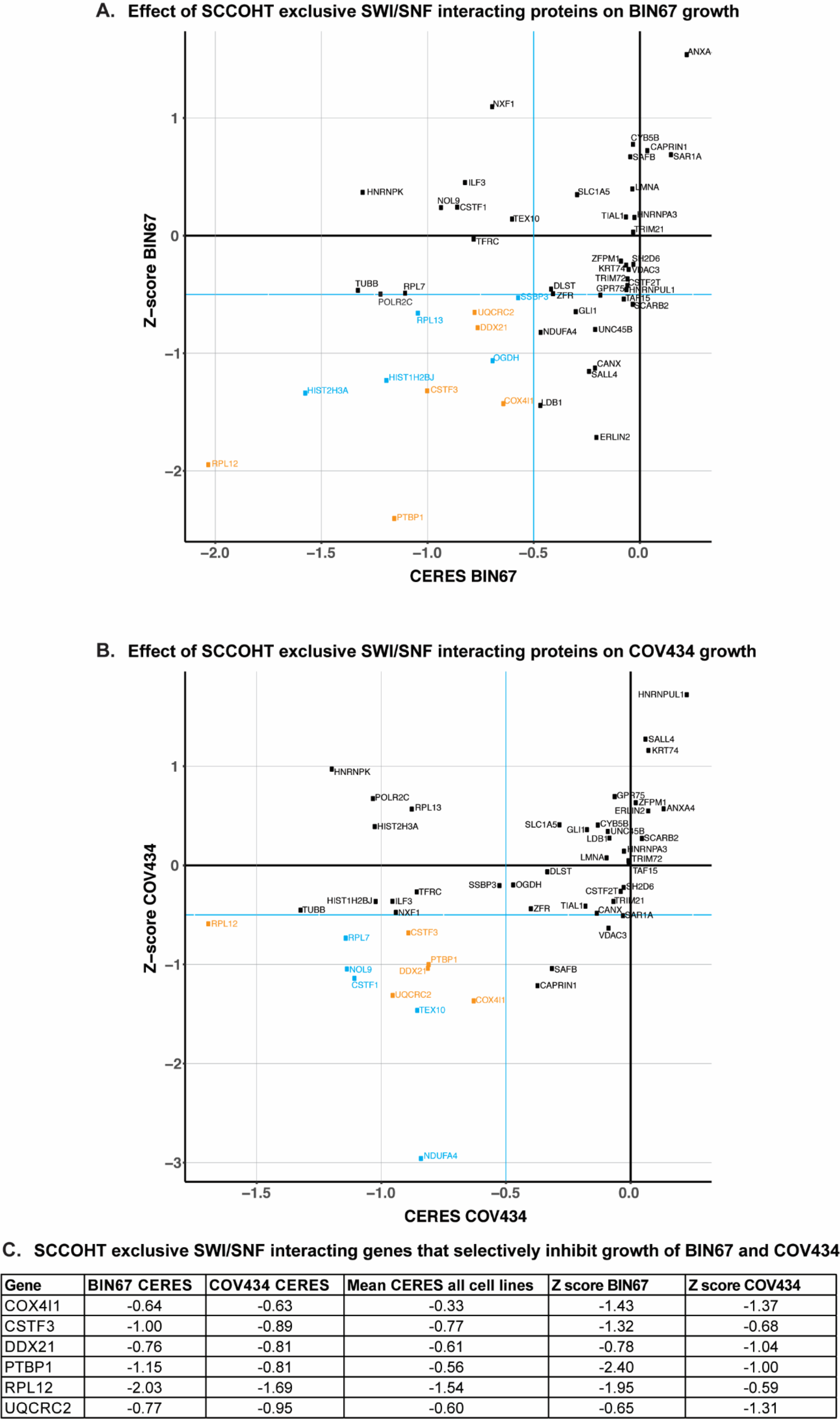
A subset of SCCOHT exclusive SWI/SNF interacting proteins provide synthetic lethal dependencies. **A-B.** Effect of CRISPR knock-out of the indicated gene on BIN67 (A) and COV434 (B) growth (CERES score) compared to the average across all cell lines (Z score) in the BROAD Avana CRISPR database. Effect on growth is reported as a CERES score, where negative scores indicate less cell growth and positive scores indicate more cell growth. A CERES score of −1.0 indicates a decrease in growth comparable to knocking out an essential gene. Genes analyzed encode the 60 proteins observed to interact with SWI/SNF exclusively in SCCOHT cell lines (Figure 7). Genes colored cyan showed a CRISPR CERES score of ≥−0.5 and a Z score of ≥−0.5 in either BIN67 or COV434. Genes colored orange showed a CRISPR CERES score of ≥−0.5 and a Z score of ≥−0.5 in both BIN67 and COV434. **C**. CRISPR CERES scores for genes that showed a CRISPR CERES score of ≥−0.5 and a Z score of ≥−0.5 in both BIN67 and COV434 as in (A-B) compared to the effect of that gene on the average growth of all cell lines.

## Discussion

### Residual SWI/SNF complex in SCCOHT lacking ATPase and PBAF-specific subunits

We have characterized the SWI/SNF complex composition in SCCOHT cells. Using IP-MS we observed that the SWI/SNF isoforms present in SCCOHT cells retain core subunit interactions, show loss of ATPase module subunits, and loss of PBAF-specific subunits. SCCOHT cells express BAF and ncBAF isoforms with reduced ATPase module subunits (Figure 1). We determined that SWI/SNF subunit association with the complex mirrors global protein expression and that loss of the PBAF-specific subunit PBRM1 is due to protein instability in the absence of SMARCA4 (Figures 2 & 3). This is consistent with the model of SWI/SNF assembly posed by Mashtalir *et al.* where PBRM1 associates with PBAF through the ATPase module^13^, which is reduced in SCCOHT. Further, a recent report by Pan *et al.* using cross-linking mass spectrometry found a loss of PBAF-specific subunits from the SCCOHT SWI/SNF complex^53^.

In all 3 SCCOHT cell lines, we detected SMARCA4-specific peptides that span parts of the protein presumably unaltered by the *SMARCA4* mutations in their respective cell line (Figure 4). It is possible that a fraction of SWI/SNF complexes exist where SMARCA4 interacts with the core subunits, and/or the SMARCA4 protein interacts with a different set of proteins. Further analysis will determine if a mixed population of SWI/SNF isoforms in SCCOHT cells appears and whether any sub-populations incorporate the mutant SMARCA4 protein. Additionally, we observed that the residual SWI/SNF complexes are necessary for SCCOHT cell viability (Figure 5), highlighting important roles for SWI/SNF non-catalytic functions in SCCOHT cells.

### Distinct PPI networks of the SCCOHT residual SWI/SNF complex

The use of probabilistic scoring to define and characterize the network of SWI/SNF interacting proteins in SCCOHT cells identified in total over 600 high confidence SWI/SNF interacting proteins in the wild-type and SCCOHT cell lines (Figure 6). The enriched GO biological process terms of translation, histone post-translational modification, telomere maintenance, viral response, chromatin organization, cell cycle progression, and RNA processing are consistent with previous IP-MS studies performed in HeLa cells bearing a wild-type SWI/SNF complex. Our data additionally identified SWI/SNF interactions with proteins involved in viral response^54^. We also found that the SCCOHT SWI/SNF complex showed interactions with less chromatin organization proteins and more RNA processing proteins compared to the wild-type SWI/SNF complex (Figure 6F). Interestingly, PBAF specific subunits did not show interactions with RNA processing proteins in the wild-type cell line; however, PBAF specific subunits did show interactions with RNA processing proteins in the SCCOHT cell lines (Figure 6), potentially indicating a change in functions of the PBAF complex in the absence of an ATPase module.

### SCCOHT residual SWI/SNF complex shows reduced interactions with chromatin organization proteins

SWI/SNF core subunits and RNA processing proteins were shared among the wild-type and SCCOHT SWI/SNF complexes (Figure 7). The wild-type SWI/SNF complex showed interactions with the NuRD chromatin remodeling and HDAC complex, consistent with previous reports that the NuRD complex interacts with SWI/SNF and shows functional antagonism in embryonic stem cells^55,56^ and in oral squamous cell carcinoma cells^57^. The absence of SCCOHT SWI/SNF interactions with NuRD warrants future investigation of potential roles of NuRD in SCCOHT cells. The wild-type SWI/SNF complex, but not the SCCOHT SWI/SNF complex, showed interactions with the enhancer associated histone methyltransferase MLL3/4 complex, which mono-methylates histone H3 lysine 4 (H3K4me1)^58^. The absence of interactions with MLL3 may point to the established role of SWI/SNF in enhancer regulation which has recently been implicated in SCCOHT^53^.

### The SCCOHT residual SWI/SNF complex has reduced interactions with nuclear pore proteins

Nuclear pore proteins and proteins involved in nuclear import interacted specifically with the wild-type SWI/SNF complex and not the SCCOHT SWI/SNF complex. The absence of nuclear pore interactions in SCCOHT suggests many functional downstream consequences on chromatin states and gene expression. Several studies have demonstrated roles for nuclear pores in establishing 3-dimensional chromatin structures referred to as topologically associated domains (TADs) and gene loops which place enhancers in close proximity to their cognate transcription start sites^59^. Taken together with SMARCA4’s recent implication in TAD regulation^60^, these results suggest that TAD structure and/or gene looping may be dysregulated due to SMARCA4 loss in SCCOHT. Enhancers and superenhancers are also known to associate with nuclear pores^59^, and consistent with their dysregulation in SCCOHT, enhancers marked by H3K4me1, acetylated histone H3 lysine 27 (H3K27ac), and MLL3/4 occupancy have been found to depend on SMARCA4 expression and activity at most sites, although a substantial number of loci showed enhancer marker occupancy dependence on SMARCA4 expression but independent of its ATPase activity^53^. Interestingly, we also observed absence of SWI/SNF PPIs with the MLL3 complex in SCCOHT cells. The SWI/SNF complex and chromatin association with nuclear pores has also been demonstrated to facilitate transcriptional memory which primes higher expression of a gene after an initial induction^61–63^. The loss of nuclear pore interactions with the SCCOHT SWI/SNF complex may suggest a loss of transcriptional memory that could affect how SCCOHT cells respond to environmental stimuli.

As several nuclear pore proteins have been shown to dynamically associate with nuclear pores and also be nucleoplasmic, the absence of nucleoporin (NUP) protein interactions we observed could be due to the nucleoplasmic pool of NUPs^64–66^. Several NUP proteins contain FG domains which may serve as low complexity domains and conceivably contribute to phase-separation of transcriptional compartments at nuclear pores^67–70^. NUP153, an FG domain containing NUP, only interacted with the wild-type SWI/SNF and may suggest dysregulation of phase separation in SCCOHT cells.

### The SCCOHT residual SWI/SNF complex has more PPIs with RNA processing proteins and TCA cycle proteins

The observations of more SWI/SNF PPIs with proteins involved in RNA processing in SCCOHT and the exclusive interaction of a subset of RNA processing proteins with the residual SCCOHT SWI/SNF complex was unexpected. As these gained RNA processing proteins appear to have the same biological functions as those shared with the wild-type complex, their effects on SCCOHT RNA processing are not obvious. However, it reflects that SCCOHT cells depend more on ribosome biogenesis, RNA splicing, and RNA 3’ end formation than other cell types. Given that we showed the residual complex has a significant impact on SCCOHT cell growth, these dependencies are likely important for SCCOHT cell viability.

The interaction of nuclear proteoforms of several tricarboxylic acid (TCA) cycle and electron transport chain (ETC) members with the SCCOHT residual SWI/SNF complex is noteworthy. Metabolic regulation is altered in many cancers providing an exciting target for drug development^71^. The SWI/SNF complex regulates tumor metabolism^72^ and mutations in SMARCA4 sensitize lung cancer cells to an oxidative phosphorylation inhibitor^73^. Recently reports have also discovered that nucleoplasmic TCA cycle proteins are involved in histone post-translational modification including histone succinylation^74^. These results warrant investigation of metabolic inhibitors for use in SCCOHT and may also indicate a role for altered histone post-translational modification in SCCOHT.

In summary, this work provides the most detailed characterization to date of the SWI/SNF PPI network in SMARCA4/A2-deficient SCCOHT cells. Furthermore, this level of analysis has provided novel functional insights in addition to identifying several biological processes that can immediately be considered for potential therapeutics. Foremost among the identified vulnerabilities in SCCOHT include central metabolism, ribosome biogenesis, and RNA processing. As the SWI/SNF complex is mutated in 20% of all cancers, these results may inform on functional consequences of SWI/SNF subunit loss in other tumor types.

## Acknowledgements

We thank SCCOHT patients and their families who inspired this work. We kindly thank Ralf Hass for the SCCOHT1 cell line and Barbara Vanderhyden for the BIN67 cell line. We thank Kevin Drenner for variant calling analysis from HAP1 whole genome sequencing data and Deon Nguyen for protein sample preparation. This work was supported by NIH R01 CA195670-01 awarded to Drs. Trent, Weissman, and Huntsman, Baylor Scott and White Research Institute award (CP7) to Drs. Hendricks, Bennett, and Spillman, and philanthropic support to the TGen Foundation to Dr. Trent.

## References

1 Kadoch, C. & Crabtree, G. R. Mammalian SWI/SNF chromatin remodeling complexes and cancer: Mechanistic insights gained from human genomics. Sci Adv 1, e1500447, doi:10.1126/sciadv.1500447 (2015).

2 Kadoch, C. et al. Proteomic and bioinformatic analysis of mammalian SWI/SNF complexes identifies extensive roles in human malignancy. Nat Genet 45, 592–601, doi:10.1038/ng.2628 (2013).

3 Stern, M., Jensen, R. & Herskowitz, I. Five SWI genes are required for expression of the HO gene in yeast. J Mol Biol 178, 853–868, doi:10.1016/0022-2836(84)90315-2 (1984).

4 Breeden, L. & Nasmyth, K. Cell cycle control of the yeast HO gene: cis- and trans-acting regulators. Cell 48, 389–397, doi:10.1016/0092-8674(87)90190-5 (1987).

5 Neigeborn, L. & Carlson, M. Genes affecting the regulation of SUC2 gene expression by glucose repression in Saccharomyces cerevisiae. Genetics 108, 845–858 (1984).

6 Cairns, B. R., Kim, Y. J., Sayre, M. H., Laurent, B. C. & Kornberg, R. D. A multisubunit complex containing the SWI1/ADR6, SWI2/SNF2, SWI3, SNF5, and SNF6 gene products isolated from yeast. Proc Natl Acad Sci U S A 91, 1950–1954, doi:10.1073/pnas.91.5.1950 (1994).

7 Peterson, C. L. & Workman, J. L. Promoter targeting and chromatin remodeling by the SWI/SNF complex. Curr Opin Genet Dev 10, 187–192 (2000).

8 Wang, W. et al. Purification and biochemical heterogeneity of the mammalian SWI-SNF complex. EMBO J 15, 5370–5382 (1996).

9 Lemon, B., Inouye, C., King, D. S. & Tjian, R. Selectivity of chromatin-remodelling cofactors for ligand-activated transcription. Nature 414, 924–928, doi:10.1038/414924a (2001).

10 Yan, Z. et al. PBAF chromatin-remodeling complex requires a novel specificity subunit, BAF200, to regulate expression of selective interferon-responsive genes. Genes Dev 19, 1662–1667, doi:10.1101/gad.1323805 (2005).

11 Alpsoy, A. & Dykhuizen, E. C. Glioma tumor suppressor candidate region gene 1 (GLTSCR1) and its paralog GLTSCR1-like form SWI/SNF chromatin remodeling subcomplexes. J Biol Chem 293, 3892–3903, doi:10.1074/jbc.RA117.001065 (2018).

12 Michel, B. C. et al. A non-canonical SWI/SNF complex is a synthetic lethal target in cancers driven by BAF complex perturbation. Nat Cell Biol 20, 1410–1420, doi:10.1038/s41556-018-0221-1 (2018).

13 Mashtalir, N. et al. Modular Organization and Assembly of SWI/SNF Family Chromatin Remodeling Complexes. Cell 175, 1272–1288 e1220, doi:10.1016/j.cell.2018.09.032 (2018).

14 Reisman, D. N., Sciarrotta, J., Wang, W., Funkhouser, W. K. & Weissman, B. E. Loss of BRG1/BRM in human lung cancer cell lines and primary lung cancers: correlation with poor prognosis. Cancer Res 63, 560–566 (2003).

15 Imielinski, M. et al. Mapping the hallmarks of lung adenocarcinoma with massively parallel sequencing. Cell 150, 1107–1120, doi:10.1016/j.cell.2012.08.029 (2012).

16 Cancer Genome Atlas Research, N. Comprehensive molecular profiling of lung adenocarcinoma. Nature 511, 543–550, doi:10.1038/nature13385 (2014).

17 Varela, I. et al. Exome sequencing identifies frequent mutation of the SWI/SNF complex gene PBRM1 in renal carcinoma. Nature 469, 539–542, doi:10.1038/nature09639 (2011).

18 Cancer Genome Atlas Research, N. Comprehensive molecular characterization of clear cell renal cell carcinoma. Nature 499, 43–49, doi:10.1038/nature12222 (2013).

19 Versteege, I. et al. Truncating mutations of hSNF5/INI1 in aggressive paediatric cancer. Nature 394, 203–206, doi:10.1038/28212 (1998).

20 Biegel, J. A. et al. Germ-line and acquired mutations of INI1 in atypical teratoid and rhabdoid tumors. Cancer Res 59, 74–79 (1999).

21 Northcott, P. A. et al. Medulloblastomics: the end of the beginning. Nat Rev Cancer 12, 818–834, doi:10.1038/nrc3410 (2012).

22 Wiegand, K. C. et al. ARID1A mutations in endometriosis-associated ovarian carcinomas. N Engl J Med 363, 1532–1543, doi:10.1056/NEJMoa1008433 (2010).

23 Jones, S. et al. Frequent mutations of chromatin remodeling gene ARID1A in ovarian clear cell carcinoma. Science 330, 228–231, doi:10.1126/science.1196333 (2010).

24 Kim, S. I. et al. Genomic landscape of ovarian clear cell carcinoma via whole exome sequencing. Gynecol Oncol 148, 375–382, doi:10.1016/j.ygyno.2017.12.005 (2018).

25 Cancer Genome Atlas Research, N. Integrated genomic analyses of ovarian carcinoma. Nature 474, 609–615, doi:10.1038/nature10166 (2011).

26 Kolin, D. L. et al. SMARCA4-deficient undifferentiated uterine sarcoma (malignant rhabdoid tumor of the uterus): a clinicopathologic entity distinct from undifferentiated carcinoma. Mod Pathol 31, 1442–1456, doi:10.1038/s41379-018-0049-z (2018).

27 Lin, D. I. et al. SMARCA4 inactivation defines a subset of undifferentiated uterine sarcomas with rhabdoid and small cell features and germline mutation association. Mod Pathol, doi:10.1038/s41379-019-0303-z (2019).

28 Ramos, P. et al. Small cell carcinoma of the ovary, hypercalcemic type, displays frequent inactivating germline and somatic mutations in SMARCA4. Nat Genet 46, 427–429, doi:10.1038/ng.2928 (2014).

29 Ramos, P. et al. Loss of the tumor suppressor SMARCA4 in small cell carcinoma of the ovary, hypercalcemic type (SCCOHT). Rare Dis 2, e967148, doi:10.4161/2167549X.2014.967148 (2014).

30 Karnezis, A. N. et al. Dual loss of the SWI/SNF complex ATPases SMARCA4/BRG1 and SMARCA2/BRM is highly sensitive and specific for small cell carcinoma of the ovary, hypercalcaemic type. J Pathol 238, 389–400, doi:10.1002/path.4633 (2016).

31 Jelinic, P. et al. Recurrent SMARCA4 mutations in small cell carcinoma of the ovary. Nat Genet 46, 424–426, doi:10.1038/ng.2922 (2014).

32 Jelinic, P. et al. Concomitant loss of SMARCA2 and SMARCA4 expression in small cell carcinoma of the ovary, hypercalcemic type. Mod Pathol 29, 60–66, doi:10.1038/modpathol.2015.129 (2016).

33 Agaimy, A., Thiel, F., Hartmann, A. & Fukunaga, M. SMARCA4-deficient undifferentiated carcinoma of the ovary (small cell carcinoma, hypercalcemic type): clinicopathologic and immunohistochemical study of 3 cases. Ann Diagn Pathol 19, 283–287, doi:10.1016/j.anndiagpath.2015.06.001 (2015).

34 Witkowski, L. et al. The influence of clinical and genetic factors on patient outcome in small cell carcinoma of the ovary, hypercalcemic type. Gynecol Oncol 141, 454–460, doi:10.1016/j.ygyno.2016.03.013 (2016).

35 Upchurch, K. S., Parker, L. M., Scully, R. E. & Krane, S. M. Differential cyclic AMP responses to calcitonin among human ovarian carcinoma cell lines: a calcitonin-responsive line derived from a rare tumor type. J Bone Miner Res 1, 299–304, doi:10.1002/jbmr.5650010309 (1986).

36 Otte, A. et al. A tumor-derived population (SCCOHT-1) as cellular model for a small cell ovarian carcinoma of the hypercalcemic type. Int J Oncol 41, 765–775, doi:10.3892/ijo.2012.1468 (2012).

37 Wang, Y. et al. Histone Deacetylase Inhibitors Synergize with Catalytic Inhibitors of EZH2 to Exhibit Antitumor Activity in Small Cell Carcinoma of the Ovary, Hypercalcemic Type. Mol Cancer Ther 17, 2767–2779, doi:10.1158/1535-7163.MCT-18-0348 (2018).

38 Carette, J. E. et al. Ebola virus entry requires the cholesterol transporter Niemann-Pick C1. Nature 477, 340–343, doi:10.1038/nature10348 (2011).

39 Lang, J. D. et al. Ponatinib Shows Potent Antitumor Activity in Small Cell Carcinoma of the Ovary Hypercalcemic Type (SCCOHT) through Multikinase Inhibition. Clin Cancer Res 24, 1932–1943, doi:10.1158/1078-0432.CCR-17-1928 (2018).

40 Shevchenko, A., Tomas, H., Havlis, J., Olsen, J. V. & Mann, M. In-gel digestion for mass spectrometric characterization of proteins and proteomes. Nat Protoc 1, 2856–2860, doi:10.1038/nprot.2006.468 (2006).

41 Rappsilber, J., Mann, M. & Ishihama, Y. Protocol for micro-purification, enrichment, pre-fractionation and storage of peptides for proteomics using StageTips. Nat Protoc 2, 1896–1906, doi:10.1038/nprot.2007.261 (2007).

42 McAlister, G. C. et al. MultiNotch MS3 enables accurate, sensitive, and multiplexed detection of differential expression across cancer cell line proteomes. Anal Chem 86, 7150–7158, doi:10.1021/ac502040v (2014).

43 Teo, G. et al. SAINTexpress: improvements and additional features in Significance Analysis of INTeractome software. J Proteomics 100, 37–43, doi:10.1016/j.jprot.2013.10.023 (2014).

44 Huntley, R. P. et al. The GOA database: gene Ontology annotation updates for 2015. Nucleic Acids Res 43, D1057–1063, doi:10.1093/nar/gku1113 (2015).

45 Doncheva, N. T., Morris, J. H., Gorodkin, J. & Jensen, L. J. Cytoscape StringApp: Network Analysis and Visualization of Proteomics Data. J Proteome Res 18, 623–632, doi:10.1021/acs.jproteome.8b00702 (2019).

46 Meyers, R. M. et al. Computational correction of copy number effect improves specificity of CRISPR-Cas9 essentiality screens in cancer cells. Nat Genet 49, 1779–1784, doi:10.1038/ng.3984 (2017).

47 Wang, Y. et al. The histone methyltransferase EZH2 is a therapeutic target in small cell carcinoma of the ovary, hypercalcaemic type. J Pathol 242, 371–383, doi:10.1002/path.4912 (2017).

48 Euskirchen, G., Auerbach, R. K. & Snyder, M. SWI/SNF chromatin-remodeling factors: multiscale analyses and diverse functions. J Biol Chem 287, 30897–30905, doi:10.1074/jbc.R111.309302 (2012).

49 Pulice, J. L. & Kadoch, C. Composition and Function of Mammalian SWI/SNF Chromatin Remodeling Complexes in Human Disease. Cold Spring Harb Symp Quant Biol 81, 53–60, doi:10.1101/sqb.2016.81.031021 (2016).

50 Arnaud, O., Le Loarer, F. & Tirode, F. BAFfling pathologies: Alterations of BAF complexes in cancer. Cancer Lett 419, 266–279, doi:10.1016/j.canlet.2018.01.046 (2018).

51 Sen, P. et al. Loss of Snf5 Induces Formation of an Aberrant SWI/SNF Complex. Cell Rep 18, 2135–2147, doi:10.1016/j.celrep.2017.02.017 (2017).

52 Szklarczyk, D. et al. STRING v10: protein-protein interaction networks, integrated over the tree of life. Nucleic Acids Res 43, D447–452, doi:10.1093/nar/gku1003 (2015).

53 Pan, J. et al. The ATPase module of mammalian SWI/SNF family complexes mediates subcomplex identity and catalytic activity-independent genomic targeting. Nat Genet 51, 618–626, doi:10.1038/s41588-019-0363-5 (2019).

54 Euskirchen, G. M. et al. Diverse roles and interactions of the SWI/SNF chromatin remodeling complex revealed using global approaches. PLoS Genet 7, e1002008, doi:10.1371/journal.pgen.1002008 (2011).

55 Yildirim, O. et al. Mbd3/NURD complex regulates expression of 5-hydroxymethylcytosine marked genes in embryonic stem cells. Cell 147, 1498–1510, doi:10.1016/j.cell.2011.11.054 (2011).

56 Hainer, S. J. & Fazzio, T. G. Regulation of Nucleosome Architecture and Factor Binding Revealed by Nuclease Footprinting of the ESC Genome. Cell Rep 13, 61–69, doi:10.1016/j.celrep.2015.08.071 (2015).

57 Mohd-Sarip, A. et al. DOC1-Dependent Recruitment of NURD Reveals Antagonism with SWI/SNF during Epithelial-Mesenchymal Transition in Oral Cancer Cells. Cell Rep 20, 61–75, doi:10.1016/j.celrep.2017.06.020 (2017).

58 Sze, C. C. & Shilatifard, A. MLL3/MLL4/COMPASS Family on Epigenetic Regulation of Enhancer Function and Cancer. Cold Spring Harb Perspect Med 6, doi:10.1101/cshperspect.a026427 (2016).

59 Pascual-Garcia, P. & Capelson, M. Nuclear pores in genome architecture and enhancer function. Curr Opin Cell Biol 58, 126–133, doi:10.1016/j.ceb.2019.04.001 (2019).

60 Barutcu, A. R. et al. SMARCA4 regulates gene expression and higher-order chromatin structure in proliferating mammary epithelial cells. Genome Res 26, 1188–1201, doi:10.1101/gr.201624.115 (2016).

61 Kundu, S., Horn, P. J. & Peterson, C. L. SWI/SNF is required for transcriptional memory at the yeast GAL gene cluster. Genes Dev 21, 997–1004, doi:10.1101/gad.1506607 (2007).

62 Brickner, J. H. Transcriptional memory at the nuclear periphery. Curr Opin Cell Biol 21, 127–133, doi:10.1016/j.ceb.2009.01.007 (2009).

63 Pascual-Garcia, P. et al. Metazoan Nuclear Pores Provide a Scaffold for Poised Genes and Mediate Induced Enhancer-Promoter Contacts. Mol Cell 66, 63–76 e66, doi:10.1016/j.molcel.2017.02.020 (2017).

64 Liang, Y., Franks, T. M., Marchetto, M. C., Gage, F. H. & Hetzer, M. W. Dynamic association of NUP98 with the human genome. PLoS Genet 9, e1003308, doi:10.1371/journal.pgen.1003308 (2013).

65 Vaquerizas, J. M. et al. Nuclear pore proteins nup153 and megator define transcriptionally active regions in the Drosophila genome. PLoS Genet 6, e1000846, doi:10.1371/journal.pgen.1000846 (2010).

66 Capelson, M. et al. Chromatin-bound nuclear pore components regulate gene expression in higher eukaryotes. Cell 140, 372–383, doi:10.1016/j.cell.2009.12.054 (2010).

67 Xu, S. & Powers, M. A. In vivo analysis of human nucleoporin repeat domain interactions. Mol Biol Cell 24, 1222–1231, doi:10.1091/mbc.E12-08-0585 (2013).

68 Milles, S. et al. Facilitated aggregation of FG nucleoporins under molecular crowding conditions. EMBO Rep 14, 178–183, doi:10.1038/embor.2012.204 (2013).

69 Frey, S. et al. Surface Properties Determining Passage Rates of Proteins through Nuclear Pores. Cell 174, 202–217 e209, doi:10.1016/j.cell.2018.05.045 (2018).

70 Kalverda, B., Pickersgill, H., Shloma, V. V. & Fornerod, M. Nucleoporins directly stimulate expression of developmental and cell-cycle genes inside the nucleoplasm. Cell 140, 360–371, doi:10.1016/j.cell.2010.01.011 (2010).

71 Anderson, N. M., Mucka, P., Kern, J. G. & Feng, H. The emerging role and targetability of the TCA cycle in cancer metabolism. Protein Cell 9, 216–237, doi:10.1007/s13238-017-0451-1 (2018).

72 Nickerson, J. A., Wu, Q. & Imbalzano, A. N. Mammalian SWI/SNF Enzymes and the Epigenetics of Tumor Cell Metabolic Reprogramming. Front Oncol 7, 49, doi:10.3389/fonc.2017.00049 (2017).

73 Lissanu Deribe, Y. et al. Mutations in the SWI/SNF complex induce a targetable dependence on oxidative phosphorylation in lung cancer. Nat Med 24, 1047–1057, doi:10.1038/s41591-018-0019-5 (2018).

74 Wang, Y. et al. KAT2A coupled with the alpha-KGDH complex acts as a histone H3 succinyltransferase. Nature 552, 273–277, doi:10.1038/nature25003 (2017).

